# Rapid genome sequencing for outbreak analysis of the emerging human fungal pathogen *Candida auris*

**DOI:** 10.1101/201343

**Authors:** Johanna Rhodes, Alireza Abdolrasouli, Rhys A. Farrer, Christina A. Cuomo, David M. Aanensen, Darius Armstrong-James, Matthew C. Fisher, Silke Schelenz

## Abstract

**Background:** *Candida auris* was first described in 2009, and has since caused nosocomial outbreaks, invasive infections and fungaemia across 11 countries in five continents. An outbreak of *C. auris* occurred in a specialised cardiothoracic London hospital between April 2015 and November 2016, which to date has been the largest outbreak reported worldwide, involving a total of 72 patients.

**Methods:** To understand the epidemiology of *C. auris* infection within this hospital, we sequenced the genomes of outbreak isolates using Oxford Nanopore Technologies and Illumina in order to type antifungal resistance alleles and to explore the outbreak within its local and global context.

**Findings:** Phylogenomic analysis placed the UK outbreak in the India/Pakistan clade, demonstrating an Asian origin. The outbreak showed similar diversity to that of the entire clade and limited local spatiotemporal clustering was observed. One isolate displayed resistance to both echinocandins and 5-flucytosine; the former was associated with a serine to tyrosine amino acid substitution in the gene *FKS1*, and the latter was associated with a phenylalanine to isoleucine substitution in the gene *FUR1*. These mutations are novel for this pathogen.

**Interpretation:** Multiple differential episodic selection of antifungal resistant genotypes has occurred within a genetically heterogenous population across this outbreak, creating a resilient pathogen and making it difficult to define local-scale patterns of transmission as well as implementing outbreak control measures.

**Funding:** Antimicrobial Research Collaborative, Imperial College London

## Introduction

The emerging fungal pathogen *Candida auris* causes nosocomial invasive infections, predominantly in intensive care units (ICU). Since its first description in 2009 in Japan (1), reports of *C. auris* infections have been reported in several countries (2–10). *C. auris* demonstrates intrinsic multidrug resistant (MDR) phenotype (9), by exhibiting high level resistance to fluconazole and varying susceptibility to other azole drugs, amphotericin B and a newly introduced class of antifungals, echinocandins (9).

In 2016, we described the first large-scale *C. auris* outbreak (April 2015 to November 2016) occurring within a single specialist cardiothoracic hospital in London (6). Due to the high uncertainty as to the time and source of introduction of *C. auris* into the hospital, the rapid development of a molecular epidemiological toolkit was required. Outbreaks of other fungal pathogens have been previously investigated using short-read whole-genome sequencing (WGS), which provided sufficient information to discriminate between isolates and their phylogenetic relationships using single nucleotide polymorphism (SNP) analysis (11–13). Recently, the handheld, portable MinION sequencer, manufactured by Oxford Nanopore Technologies, UK (ONT) has made rapid WGS widely available in the field, and has been successfully used to analyse the molecular epidemiology of recent *Salmonella*, and Ebola and Zika viruses outbreaks (14–16).

Here, we describe the first use of the MinION nanopore sequencing technology to determine the genetic epidemiology of this fungal outbreak, both within the UK hospital and also within a global context, alongside reannotating the genome of *C. auris* and defining novel antifungal resistance alleles.

## Methods

### Fungal isolates

Twenty-eight *C. auris* isolates were studied, consisting of 26 clinical isolates from 22 patients and two isolates collected from the room of a patient known to be colonised with *C. auris* in the ICU (Table 1). The identification of *C. auris* was conducted by Matrix-assisted laser desorption/ionization Time of Flight (MALDI-TOF) mass spectrometry (Brucker Daltonics, Fremont, CA, USA) using a formic acid – acetonitrile extraction procedure. Scores were interpreted as >2.00 for species-level identification.

**Table 1:**
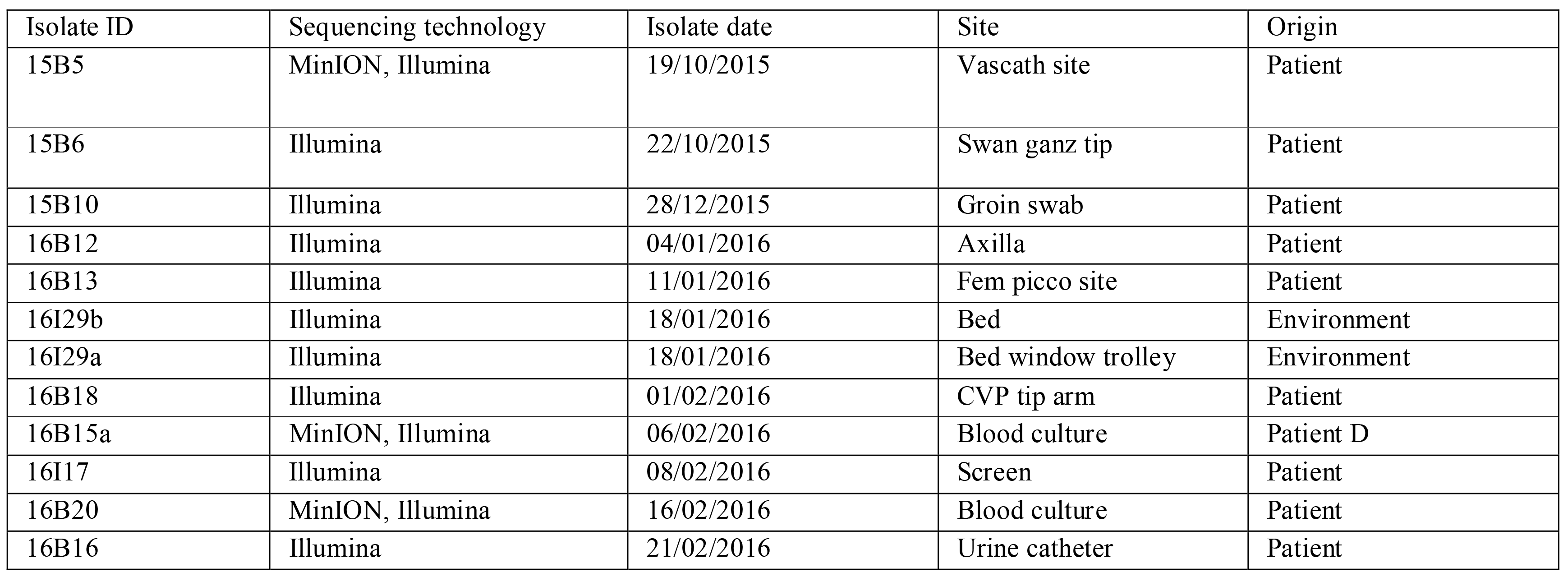
Clinical isolates of *C. auris* included in this study

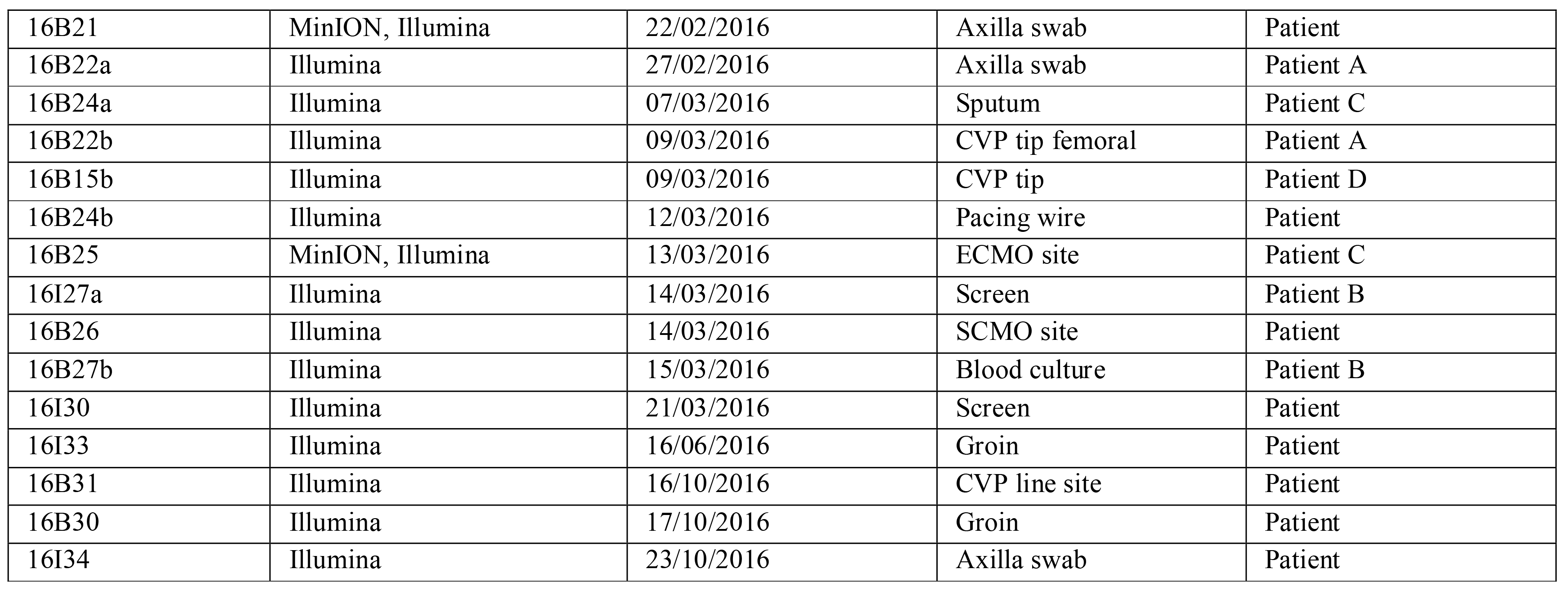

**Table 2:**
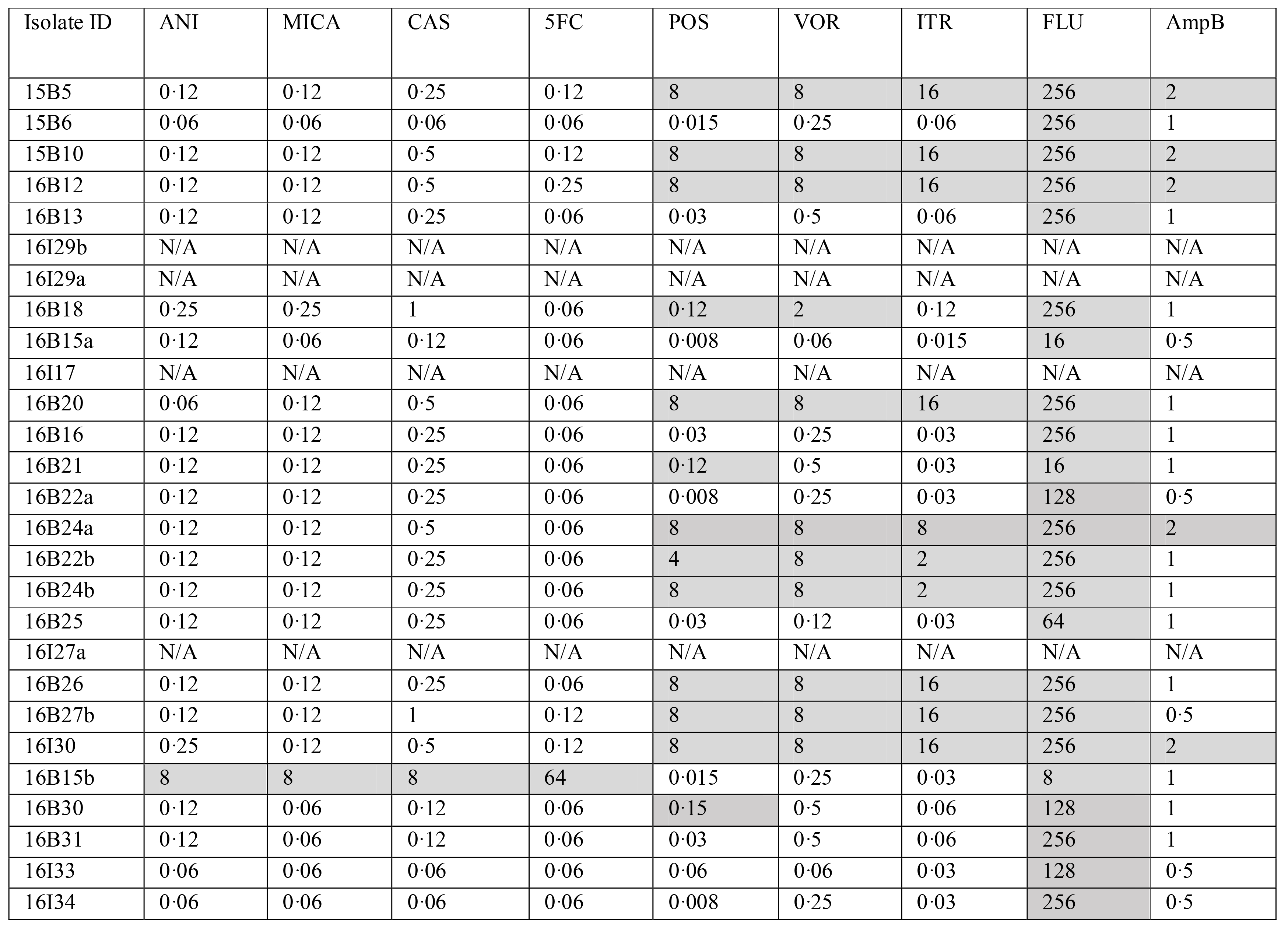
MICs at time of isolation of all isolates included in this study. MICs deemed above these breakpoint values (based on *Candida albicans* CSLI), and therefore resistant, are shaded in grey. ANI = Anidulafungin; MICA = micafungin; CAS=Caspofungin; 5FC=5-fluorocytosine; POS = posaconazole; VOR = voriconazole; ITR = itraconazole; FLU = fluconazole; AmpB = Amphotericin B. N/A = No MICs carried out.

### Library preparation and sequencing

Five clinical isolates (Table 1) representing a 146-day time frame were chosen for sequencing using the hand-held MinION sequencer (Oxford Nanopore Technologies, Oxford, UK).

Twenty-four isolates (Table 1), including the five clinical isolates sequenced using MinION, were chosen for Illumina sequencing using Nextera library preparation method as part of the MicrobesNG service (University of Birmingham, UK). These isolates represented a 155-day time frame, and included isolates from patients and the environment around infected patients. Isolates of *C. auris* for both MinION and Illumina sequencing were cultured as described in Supplementary Methods. All raw reads in this study have been submitted to the European Nucleotide Archive under the project accession PRJEB20230. Details of genome assembly and annotation are described in Supplementary Methods.

### Alignment of Illumina reads and phylogenetic analysis

Raw Illumina reads were quality checked using FastQC (v0.11.3; Babraham Institute), and trimmed using Trimmomatic (v0.30) based on a Phred quality score of 15. Alignment of reads to the 16B25 reference genome and variant calling were carried out as described in Rhodes and colleagues (17).

Phylogenies for whole genome SNP data were constructed and visualised as described in Rhodes and colleagues (17). The rate of evolution (represented as the number of substitutions per day) along the tree topology was estimated using TempEst v1.5 (18), calibrated with sampling times. Root-to-tip regression was calculated and the root of the tree was selected to maximise *R*^2^.

### Mutation identification in *ERG11*, *FKS1* and *FUR1* genes

Orthologous sequences to *C. albicans ERG11* (SC5314) were extracted from each *C. auris* genome. Sequences were evaluated for amino acid substitutions to mutations within hot spot regions in *C. albicans* (19) as described in Lockhart and colleagues (9). Predicted *FKS1* and *FUR1* genes from the genome annotation were used to identify presence or absence of mutations in *C. auris* isolates.

## Results

We sequenced 25 clinical *C. auris* isolates from a recently described outbreak (6), along with two environmental samples to better represent the overall genetic diversity within the impacted hospital. We also sequenced eight isolates derived from four patients taken days apart to establish possible within-patient diversity.

### Rapid generation of outbreak-specific *C. auris* reference genome and Illumina sequencing

We assembled five high quality hybrid *de novo* reference genomes for *C. auris* using Illumina short-read sequences and MinION long-read sequences rapidly generated over 48 hours. Five isolates (15B5, 16B21, 16B25, 16B20 and 16B15a) were chosen to cover a range of dates (October 2015 to March 2016). Isolate 16B25 had the best overall assembly quality of 110 contigs, N_50_ = 396,317 bp and an estimated genome size of 12–3 Mb (Table 3 and Supplementary Table S1). 98.94% of the 16B25 assembly mapped to the *C. auris* genome B8441 assembled by Lockhart and colleagues (9).

**Table 3:**
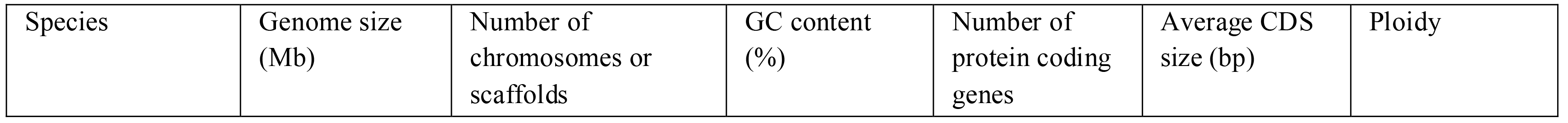
Summary of assembly and annotation statistics of the *C. auris* 16B25 genome, the *C. auris* B8441 reference (9), the *C. auris* Ci 6684 reference (21), and other pathogenic *Candida* species reference genomes(20)

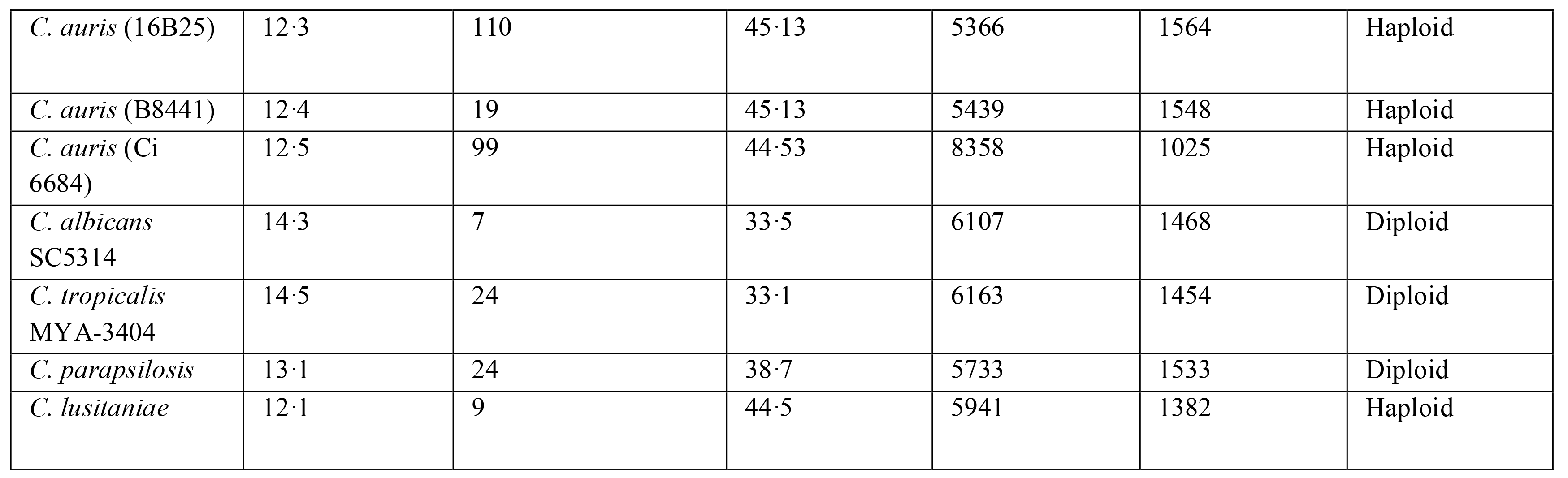

We generated an average of 5·2 million Illumina reads passing quality control for 27 isolates recovered during the outbreak that mapped closely (average 95·5%) to our reference genome (Supplemental Material Table S2). The rapid availability of long reads from MinION sequencing demonstrates this technology is ideally used in an outbreak setting for providing high-quality contiguous assemblies. A total of 5,366 protein-coding genes, 4 rRNAs and 156 tRNAs were predicted using the genome annotation pipeline described in Supplementary Methods. Table 3 summarises the general features of 16B25, along with other pathogenic *Candida* genomes. The number of protein coding genes presented here is in line with the predicted number of genes in *C. lusitaniae* (20) (*n* = 5,941), the closest known relative of *C. auris*.

There are fewer protein coding genes, tRNAs and rRNAs predicted in this genome than previously reported for *C. auris* Ci 6684 in Chatterjee and colleagues (21), as shown in Table 3. Running our annotation pipeline on the B8441 isolate presented in Lockhart and colleagues (9) found similar numbers of protein coding genes, rRNAs and tRNAs (Table 3). Therefore, the different total numbers between 16B25 and Ci 6684 are likely due to the different annotation pipelines and not the quality of the reference assemblies. The number of protein coding genes identified in Chatterjee and colleagues is likely inflated due to over-prediction of short sequences, lack of filtering of repetitive sequences, and using only GenemarkS to predict the start of genes; our pipeline used additional criteria to achieve a predicted set of high-confidence genes.

### Phylogenetic analysis reveals an Indian/Pakistani origin of *C. auris* outbreak

Phylogenetic analysis based on whole genome SNPs revealed the UK outbreak had an Indian/Pakistani origin (Figure 1). SNP calls for isolates from Venezuela, India, Pakistan, Japan and South Africa (9) were also included to add geographic context to the outbreak. The UK outbreak isolates were in the same clade as those from India and Pakistan (Figure 1a); on average, 240 SNPs separated UK outbreak isolates from isolates collected in India and Pakistan. There were no known patient travel links to India or Pakistan prior to admission into hospital, however. We found an average of 84 SNPs separating isolates within the UK outbreak; later isolates exhibited only 55 SNPs between them (October 2016), compared to earlier isolates (January 2016) that showed an average 130 SNPs separating them.

**Figure 1.**
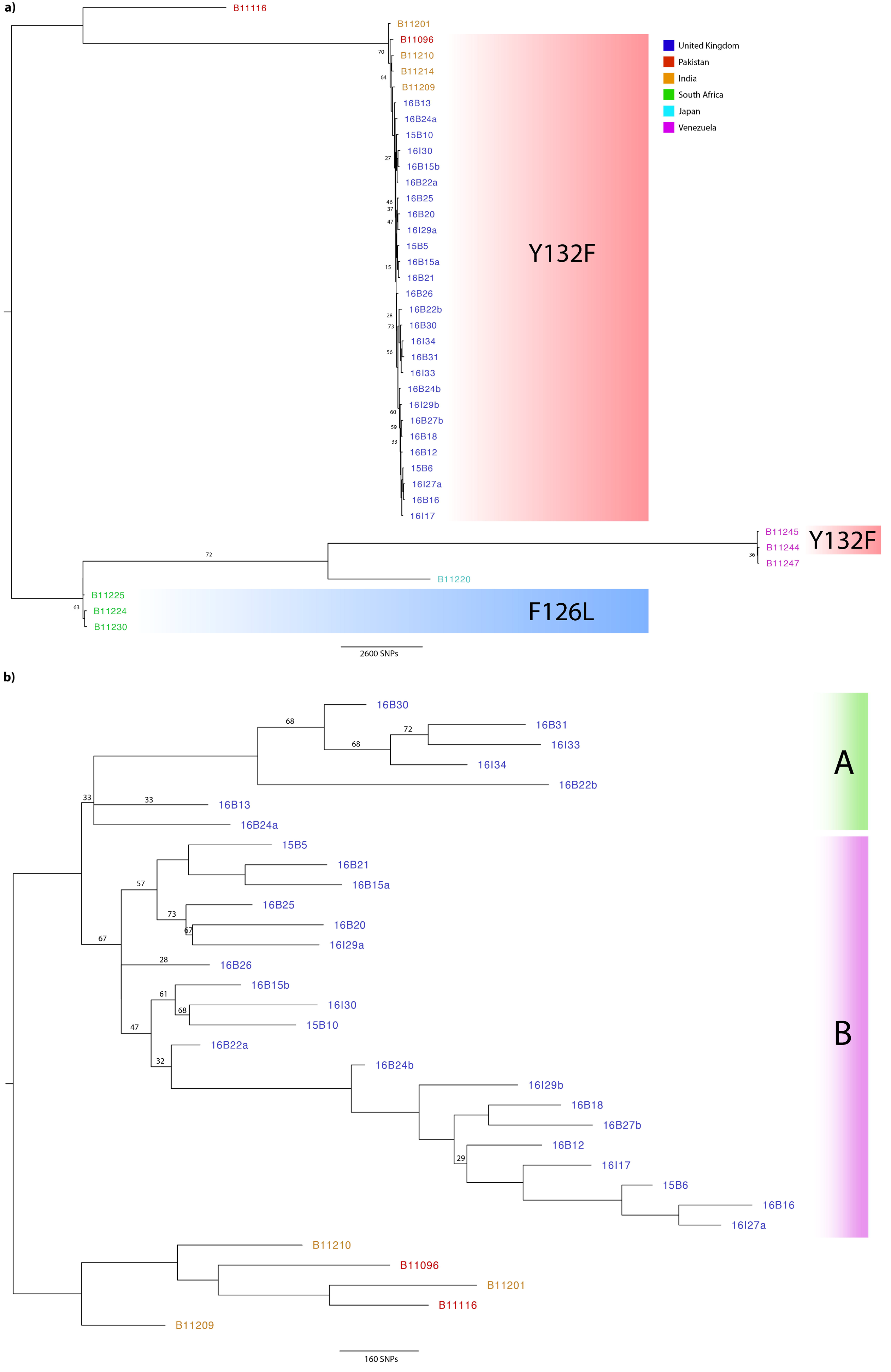
Phylogenetic analysis of *C. auris* isolates with bootstrap support (500 replicates) performed on WGS SNP data to generate unrooted maximum-likelihood phylogenies. Branches supported to 75%or higher unless otherwise stated. Branch lengths represent the average expected rates of substitution per site. **a)** Outbreak isolates from the UK (shown in blue) were combined with isolates from around the globe, including India (orange), Pakistan (red), Venezuela (pink), Japan (turquoise), and South Africa (green), to infer a possible geographical origin. Isolates with known mutations in the *ERG11* gene associated with resistance to fluconazole in *C. albicans* are shaded: Y132F as red, and F126L as blue **b)** Given the likely Indian/Pakistani origin of the outbreak isolates, phylogenetic analysis was repeated (as stated above), excluding isolates from South Africa, Venezuela and Japan to illustrate the UK outbreak. Isolates separating either into Cluster A (green) or B (purple) are depicted to infer likely introductions into the hospital.

Fitting root-to-tip regression showed there was a linear relationship between sampling time (days) and the expected number of nucleotide substitutions along the tree, demonstrating clock-like evolution across the time-scale of the outbreak (Figure 2). The evolutionary rate of nuclear DNA (calculated from the slope of the regression) equated to 1.5204^e−3^ substitutions per site, comparable with nuclear DNA of other fungal species such as *Schizosaccharomyces pombe* beer strains (3.0^e−3^ (22)) and *Saccharyomyces cerevisiae* (5.7^e−3^ (23)). The time to the most recent common ancestor (TMRCA) was estimated to be early March 2015, one month prior to the first patient identified with a *C. auris* infection. One isolate, 16B22b, was identified as being less diverged than average for the sampling date; five isolates (16B15b, 16B26, 16B25, 16B22a and 16B24a) were identified as being more diverged than average for the sampling date given.

**Figure 2.**
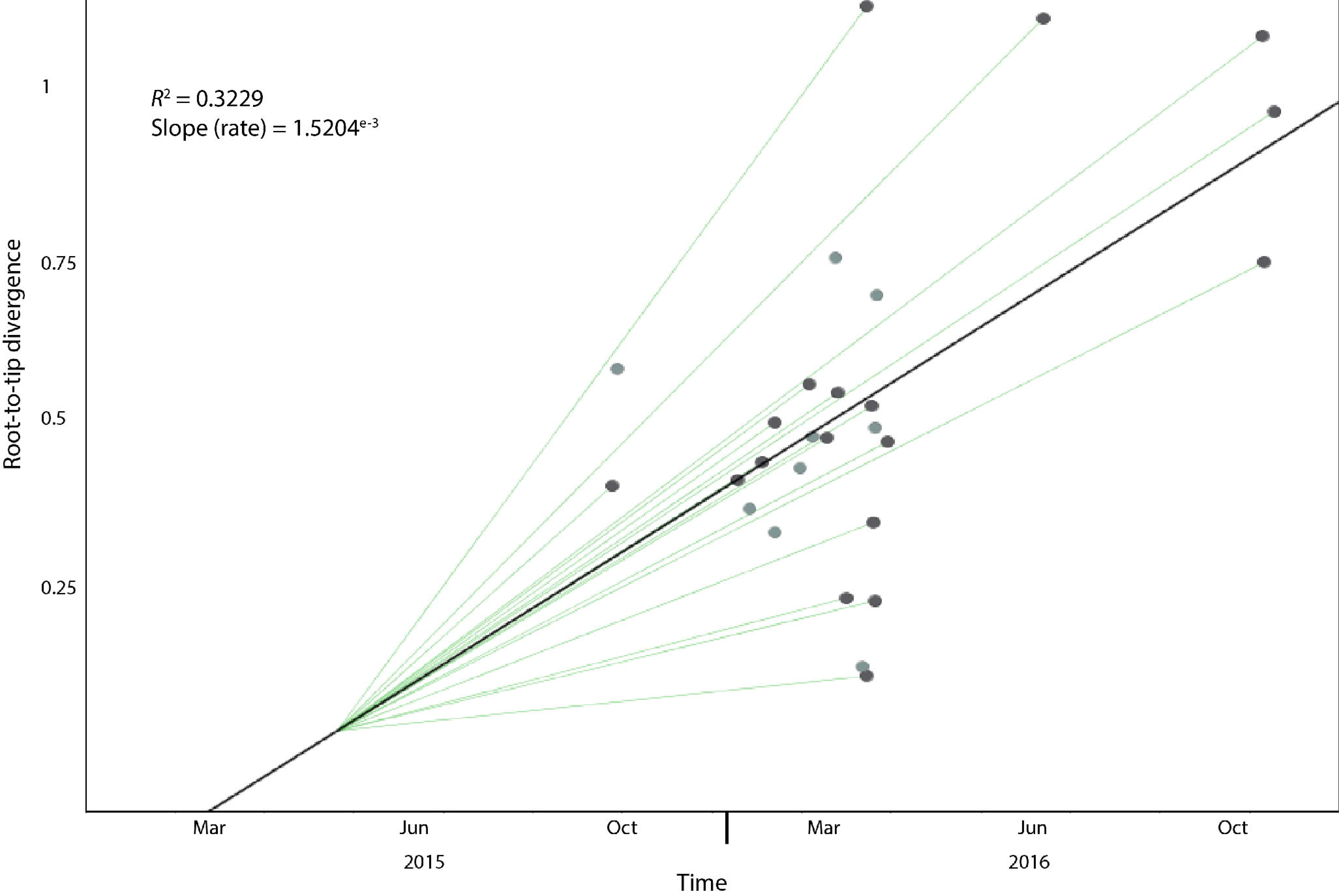
Root-to-tip regression analysis of all 28 *C. auris* outbreak isolates. Genetic distance is plotted against sampling time for the phylogeny of the *C. auris* outbreak. Each data point represents a tip on the phylogeny.The *R*^2^ for the regression and the slope, reflecting the evolutionary rate (in substitutions per site per day) is also shown.

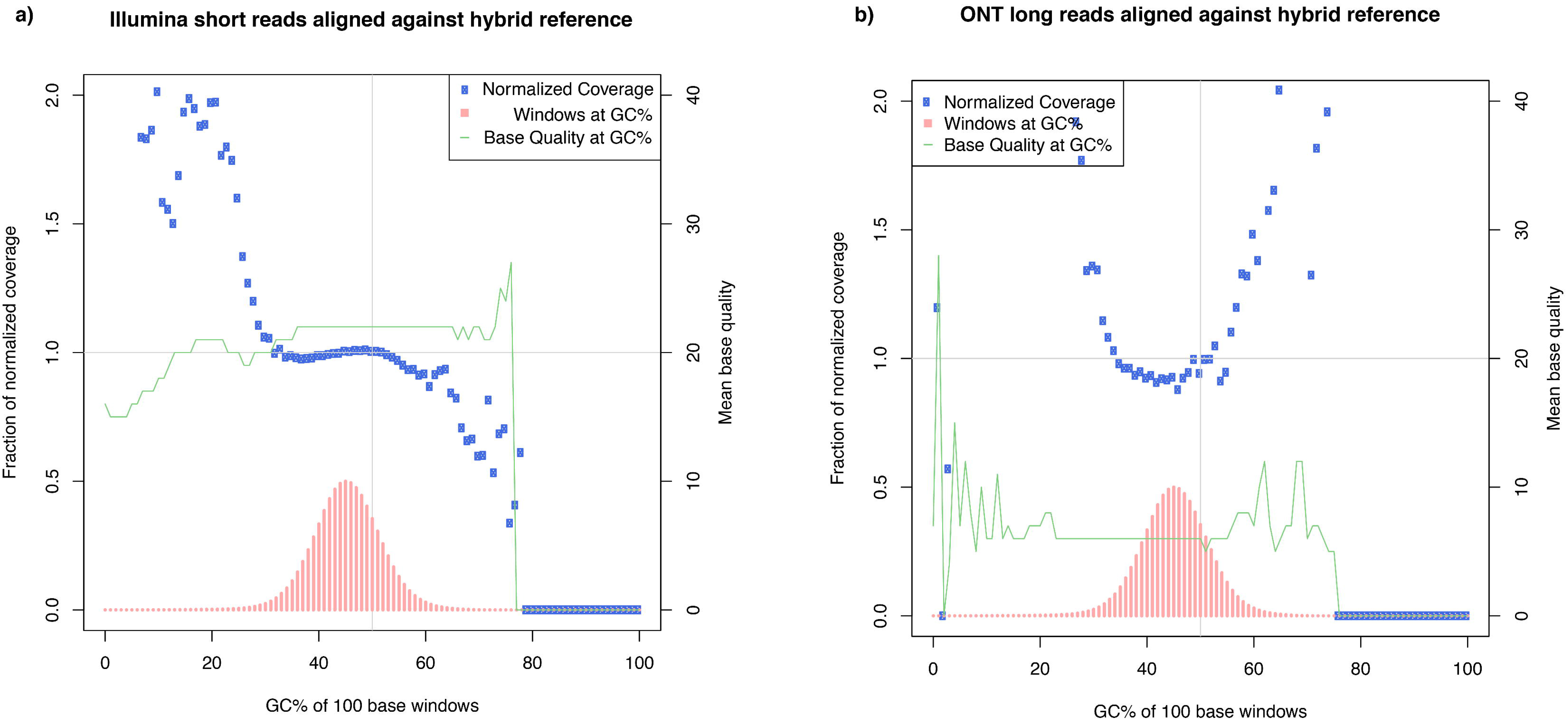

### Clinical isolates of *C. auris* show multi-drug resistance

Overall, 14 isolates displayed MDR to two or more classes of antifungal drugs. Only five isolates displayed resistance to one drug, fluconazole. All isolates expressed elevated levels of resistance to fluconazole(MIC: >256 ug/ml), with varying levels of resistance to itraconazole (MIC: 0.03 ug/ml - >16 ug/ml), voriconazole (MIC: 0.12 ug/ml - 8 ug/ml) and posaconazole (which has not been previously reported in *C. auris)*. Four isolates also displayed resistance to amphotericin B (MIC: >2 ug/ml).

One isolate (16B15b) displayed elevated levels of resistance to all echinocandins (MIC: >8 ug/ml), but remained susceptible to all azole drugs, with the exception to fluconazole. 16B15b also displayed high levels of resistance to flucytosine (MIC >64 ug/ml), which was not seen in the other isolates; therefore, both mutations and associated resistance were unique to this outbreak isolate. This isolate belonged to a patient who received anidulafungin for 7 days for pancolitis, developing *C. auris* candidaemia 11 days afterwards, at which point isolate 16B15a was recovered. Treatment was switched to amphotericin and 5-flucytosine for 2 weeks. Six days after completing this treatment, a pan-resistant *C. auris* (16B15b) was recovered from the vascular tip. One non-synonymous SNP (nsSNP), causing a serine to tyrosine substitution (S652Y) was identified in the *C. auris FKS1* gene; a similar mutation (S645Y) in *FKS1* has been associated with echinocandin resistance in *C. albicans* (24). Another nsSNP caused a phenylalanine to isoleucine substitution (F211I) in *FUR1*, which has a role in 5-flucytosine resistance (25). Neither of these mutations have been reported previously.

Orthologous sequences to *C. albicans ERG11* were screened for substitutions that conferred known fluconazole resistance mutations (19). The outbreak isolates all had the Y132F substitution in *ERG11*, confirming an Indian/Pakistani origin. Lockhart and colleagues also found that these substitutions were strongly correlated with geographic clades (9).

### Interpretation of typing results in relation to epidemiology of the outbreak

*C. auris* outbreak isolates grouped into two phylogenetic clusters (A and B; 26% and 74% respectively) (Figure 1b). Cluster A comprised seven isolates from 2016 that had on average 245 SNPs that distinguished it from Cluster B. On average 130 SNPs separated isolates within Cluster B. Cluster A was introduced into the hospital in early 2016 and formed the dominant outbreak strains towards the end of the outbreak, with only 55 SNPs separating those isolates. The phenotypic antifungal resistance varied among these isolates: all expressed high level fluconazole resistance (MIC: 128–256 ug/ml) and susceptibility to echinocandins (micafungin, caspofungin and anidulafungin MIC: 0.06–0.12 ug/ml) and 5-flucytosine (MIC 0.06 ug/ml), two clinical isolates expressed cross azole resistance (MIC: 4–8 ug/ml posaconazole, 8 ug/ml voriconazole, 2–16 ug/ml itraconazole). Phenotypic antifungal susceptibility profiling may therefore not be used reliably for addressing genetically indistinguishable strains during nosocomial transmission analysis.

Three patients with isolates in Cluster A were admitted to the ICU. Two patients acquired *C. auris* whilst staying in ICU, and another patient acquired *C. auris* on a surgical admission unit geographically placed next door to the high dependency unit in the same month, but not overlapping with the ICU patients. Transmission of *C. auris* between different units within this hospital likely occurred *via* the movement of *C. auris*-positive patients or contaminated equipment. However, because we did not sequence isolates from all patients during the outbreak alongside the heterogeneous nature of the founding population, we are currently unable to establish routes of transmission in more detail.

### Analysis and interpretation of sequencing result in relation to epidemiology of individual patient transmission during the outbreak

Eight isolates within this study were sequential pairs of isolates from four separate patients (Table 1); we hypothesised that there may have been nosocomial horizontal transmission between patients and/or their surrounding environment, as suggested in previous studies (2,3,5,26), for the following pairs of isolates: 16B22a and 16B22b from patient A (isolated 12 days apart); 16I27a (MICs were not carried out for this isolate) and 16B27b from patient B (isolated 1 day later); 16B24a and 16B24b from patient C (isolated 5 days apart); and 16B15a and 16B15b from patient D (isolated 32 days apart).

In patient A, 16B22a (recovered from the axilla) showed resistance to fluconazole only (Table 2). The subsequent isolate from this patient, 16B22b (isolated from a central line tip) exhibited resistance to all azole drugs (Table 2). A large number of SNPs (277 SNPs) separated the two isolates (Figure 1), which also separated into different phylogenetic clusters (16B22a in Cluster B and 16B22b in Cluster A) suggesting independent acquisition of infection by this patient within the unit. The two isolates from patient B similarly differed by a large number of SNPs (164 SNPs), but were both placed in Cluster B. Given that 16I27a (from a body screen sample) was isolated one day prior to 16B27b (recovered from a positive blood culture), it is likely that this patients diversity represents an heterogenous infecting population. In patient C, 16B24b was isolated from a clinical pacing wire sampled five days after the initial isolation of 16B24a from sputum sample. These two isolates differ by over 400 SNPs, shown by their disparate position in the phylogeny (Figure 1) with 16B24a placed in Cluster A, and 16B24b placed in Cluster B. 16B15b showed raised MICs to all echinocandins and flucytosine, which was not displayed in 16B15a. These two isolates were separated by only 120 SNPs, and were phylogenetically placed in Cluster B (Figure 1b). Our results suggest that none of these patients were infected with a single, clonally propagating *C. auris* strain, and instead the hospital (and all patients) was seeded with a genetically heterogenous population.

Three isolates (16B25, 16B20,16I29a) clustered closely together with only 99 SNPs difference. 16I29a was recovered from the environment around a *C. auris*-positive patient (the isolate was not sequenced). The patient from which 16B20 was recovered was present in an adjacent side room at the same time, suggesting a potential transmission between these patients. However, it remains unclear how the organism may have been transferred between the two rooms and patients. The third isolate of this cluster (16B25) was recovered from a patient present in the same room where 16I29a was recovered. The patient was placed in this room 22 days after the previous *C. auris*-positive patient had left the room, and became positive within 14 days of being in this isolation room.

Isolates 15B5, 16B15b and 16B21 are phylogenetically related (separated by 95 SNPs; Figure 1b), and were recovered from patients sharing the same bay. The bay was initially populated with the patient from whom isolates 15B5 and 16B15b were recovered, and then saw the introduction of the patient from which 16B21 was recovered. Both examples of spatiotemporal clustering suggest that there may be environmental persistence of *C. auris* resulting in transmission to patients.

## Discussion

*C. auris* is an MDR fungal pathogen, capable of causing invasive infections. Here we report WGS of *C. auris* infections from the largest outbreak to date, occuring between April 2015 and November 2016 in a London hospital ICU which spread to two other wards.

A gold standard reference genome for the outbreak was assembled using long MinION-generated reads, and Illumina short reads. Whilst the GC bias and base quality in >80% GC regions of the genome was similar in both Illumina and MinION sequencing, Illumina was more consistent across the whole genome (Figure S1). MinION reads displayed wide variation in base quality in >85% AT regions, ranging from a quality score of zero to 1.4, whereas Illumina reads ranged between quality scores of 0·75 and 0·85 for regions >85% AT. ONT has since released new chemistry that improves read quality, and therefore the variant calling, which will provide a competitive alternative to Illumina sequencing in both outbreak settings and routine research.

The speed of the MinION sequencer allowed rapid assembly and mapping of Illumina short reads to call high-confidence SNPs, which is of great importance in an outbreak (14,15). SNPs in *ERG11* correlated with known *C. albicans* hotspots (19) conferring resistance to the frontline drug fluconazole. All *C. auris* isolates in this study that exhibited high MICs to fluconazole (>250 ug/ml) contained amino acid substitutions known to cause resistance to fluconazole (19). Fourteen isolates were MDR, posing an important clinical challenge in the treatment of *C. auris* infections. Within this study we have also highlighted resistance to posaconazole, which has not previously been reported in *C. auris*, alongside echinocandin and flucytosine resistance in one MDR isolate.

Echinocandin resistance is linked to mutations in the *FKS* genes in other *Candida* species (27), and our analysis identified the S645Y mutation in *FKS1* in an echinocandin-resistant isolate. Flucytosine resistance was observed in the same isolate, and was associated with the F211I mutation in *FUR1*. Our analysis suggests that systemic echinocandin and flucytosine treatment can rapidly select for resistant genotypes across the outbreak timescale. Rapid MinION sequencing of *C. auris* isolates would allow drug resistance mutation identification, providing time-sensitive information in a clinical setting.

Phylogenomic analysis showed weak support for monophyletic status of isolates within the hospital, suggesting multiple introductions of *C. auris* occurred. On average, only 240 SNPs separated the UK outbreak isolates from the Indian/Pakistani clade, clearly showing the UK outbreak had an Asian origin. When compared to other sequenced Asian *C. auris* isolates (Table S3) in Lockhart and colleagues (9), these SNP numbers suggest an anomalous amount of diversity within this outbreak. Future alignments will require clade-specific references due to the large evolutionary distances between South America/Africa and the Indian/Pakistani clades. Although the mode of introduction into the UK is unknown, temporal analysis of the outbreak isolates placed the most recent common ancestor as early March 2015, which correlates closely to the first confirmed infection within the hospital one month after this, suggesting a recent introduction into the UK.

Only one third of SNPs were shared between hospital environment isolates (bed and trolley of a confirmed *C. auris* infected patient). *C. auris* was also recovered from inanimate surfaces, suggesting a population of genotypes are capable of contaminating the hospital environment, causing onward infection of human hosts (9). Given SNP differences between isolates recovered from the same patient at different bodily locations (Patients A, B and C), either the outbreak diversity is due to multiple introductions, which is unlikely as all patients screened negative for *C. auris* upon admission, or a genetically heterogenous population seeded the hospital. Further, given the substantial number of SNPs separating the two isolates that were recovered from Patient C 24 hours apart, it appears clear that genetically disparate isolates are capable of infecting the same host at the same time. The genomic diversity of *C. auris* within this outbreak makes mapping local-scale transmission events difficult as genetic bottlenecks may result in rapid changes in allele frequencies within local spatiotemporal scales. Clearly, sequencing multiple isolates of *C. auris* from patients alongside those from other UK outbreak settings is needed to more finely resolve the population genomic structure of this pathogen in order to understand transmission dynamics.

This study represents the first use of ONT MinION sequencer on a human fungal pathogen. The association of nsSNPs in *FKS1* and *FUR1* with echinocandin and flucytosine resistance respectively are both clinically relevant and novel. Further investigation into these mutations is required to confirm these associations. Epidemiological analysis suggests that contact tracing was not sufficient to resolve fine-scale spatiotemporal processes across the outbreak due to the multiple differential episodic selection occurring across a genetically heterogenous *C. auris* population. Future research into *C. auris* should focus on sequencing many isolates from the same patient, from multiple body sites, in order to correctly establish the nature of persistence and transmission of *C. auris* within hospital environments; the genomic approaches underpinning this study will likely be cornerstones of future research into this increasingly important infectious disease.

## Acknowledgements

We thank Oxford Nanopore Technologies for their generous contribution of flow cells and nanopore sequencing kits. We also wish to extend our thanks to Nicholas J. Loman (University of Birmingham) for providing insight into the initial conception of experiments, and to Anastasia Litvintseva, Nancy A. M. Chow and Kizee Etienne (Centers for Disease Control, USA) for the provision of additional sequence data. We also thank the sequencing team at Wellcome Trust Sanger Institute for sequencing four of the isolates presented here.

J.R. was supported by an Antimicrobial Research Collaborative (ARC) early career research fellowship(RSRO_54990).

R.A.F. was supported by an MIT/Wellcome Trust Fellowship.

M. C. F. was supported by the Natural Environmental Research Council.

D. A-J was supported by NERC and the Wellcome Trust.

C. A. C was supported by the National Institute of Allergy and Infectious Diseases, National Institutes of Health, Department of Health and Human Services, under grant number U19AI110818.

D. M. A. supported by the Wellcome Trust (Grant no. 099202)

Genome sequencing was provided by MicrobesNG (http://www.microbesng.uk), which is supported by the BBSRC (grant number BB/L024209/1)

## Author contributions

Clinical and outbreak data analysis: S.S.

Collected isolates: S.S.

Conceived experiments: J.R., M.C.F., D.A-J., S.S.

DNA extractions: A.A., J.R.

MinION DNA sequencing: J.R.

Illumina sequencing: D.M.A. and MicrobesNG

Bioinformatic analysis: J.R.

Genome annotation: R.A.F., C.A.C.

Manuscript preparation: J.R., R.A.F., A.A, S.S.

